# Evolution and origin of sliding clamp in bacteria, archaea and eukarya

**DOI:** 10.1101/2020.10.09.332825

**Authors:** Sandesh Acharya, Amol Dahal, Hitesh Kumar Bhattarai

## Abstract

Replication of DNA is an essential process in all domains of life. A protein often involved without exception in replication is the sliding clamp. The sliding clamp encircles the DNA and helps replicative polymerase stay attached to the replication machinery increasing the processivity of the polymerase. In eukaryotes and archaea the sliding clamp is called the Proliferating Cell Nuclear Antigen (PCNA) and consists of two domains. This PCNA forms a trimer encircling the DNA as a hexamer. In bacteria, the structure of the sliding clamp is highly conserved, but the protein itself, called beta clamp, contains three domains, which dimerize to form a hexamer. The bulk of literature touts a conservation of the structure of the sliding clamp, but fails to recognize conservation of protein sequence among sliding clamps. In this paper we have used PSI blast to the second interation in NCBI to show a statistically significant sequence homology between *Pyrococcus furiosus* PCNA and *Kallipyga gabonensis* beta clamp. The last two domains of beta clamp align with the two domains of PCNA. This homology data demonstrates that PCNA and beta clamp arose from a common ancestor. In this paper, we have further used beta clamp and PCNA sequences from diverse bacteria, archaea and eukarya to build maximum likelihood phylogenetic tree. Most, but not all, species in different domains of life harbor one sliding clamp from vertical inheritance. Some of these species that have two or more sliding clamps have acquired them from gene duplication or horizontal gene transfer events.

## Introduction

Replication, transcription and translation are fundamental events occurring in all living organisms. Homologous machineries exist for the transcription and translation pathways of all cells however replication machineries vary considerably among the different domains, mainly between bacteria and archeae/eukarya(Yao & O’Donnell 2016)⍰. DNA polymerases, primases and helicases involved in replication process are quite distinct and non-homologous between prokaryotes and eukaryotes(Yao & O’Donnell 2016)⍰. Interestingly two replication proteins, DNA sliding clamps and clamp loader share structural homology in the different domains of life.

DNA sliding clamp proteins, found in all organisms are essential components of DNA polymerase enzyme as they bind the enzyme and prevent it from dissociating from the template DNA strand. The sliding clamp proteins are called beta clamp in bacteria and proliferating cell nuclear antigen (PCNA) in eukaryotes(Bruck & O’Donnell 2001)⍰. Since the replication machinery of archaea demonstrates similarity to eukaryotic replication machinery than to bacterial replication machinery, archaea contain PCNA and not beta clamp. While beta clamp is a homo-dimer consisting two sub-units of three domains(Kong et al. 1992)^⍰^, PCNA is a homo-trimer made up of three subunits of two domains(Matsumiya 2001)⍰. In some archaea of Crenarchaeal sub-domain, three PCNA homologs are present, which exist as monomers in solution and self-assemble in to a functional hetero-trimer(Dionne et al. 2003; Williams et al. 2006)^⍰^. Despite the difference in the sub-units and domains, all sliding clamps assemble to form a ring-shaped protein, such that the ring is wide enough to accommodate double-stranded DNA(Kelman 1995; Jeruzalmi et al. 2002)⍰. The architecture consists of outer negatively charged surface of continuous beta-sheets, with positively charged inner surface composed of alpha helices(Kelman 1995; Hedglin et al. 2013)^⍰^. In spite of sub-unit compositions, the 3-D structure of the sliding clamp has been highly conserved throughout evolution.

Sliding clamps encircle DNA(Stukenberg et al. 1991)⍰ and thereby, physically tether the DNA polymerase to the DNA template, hence increasing the processivity of the enzyme(Stukenberg et al. 1991)⍰. Previously, the function of sliding clamps was thought to be limited only to maintain the processivity of DNA polymerase. However, with recent studies, various interactions of sliding clamps with other proteins have been recognized that shed light on different roles of sliding clamps in the cell. PCNA has been known to interact with several proteins like XPG, MSH3, MSH6, MCH1, PMS2, hMYH, Fen1 endonuclease, and DNA ligase I suggesting roles in nucleotide excision repair, mismatch repair, base excision repair, and maturation of Okazaki fragments(Parker et al. 2001)⍰. Similarly, the prokaryotic homolog of PCNA, viz. beta-clamp, has been observed to interact with MutS, PolI, DNA ligase, PolII and PolV suggesting roles in mismatch repair, processing of Okazaki fragments, and DNA repair(Wilkins 2000; López de Saro & O’Donnell 2001)^⍰^. Thus, not only the 3-D structure, but also interactions and functions are conserved between sliding clamps of prokaryotes and eukaryotes.

The existing body of information reveals that bacterial beta clamps and eukaryotic PCNA share structural homology. It has been reported that beta clamps and PCNA share no sequence homology at all although beta clamp proteins and PCNA are highly conserved in bacteria and in archaea/eukaryotes, respectively(Kong et al. 1992; Bruck & O’Donnell 2001; Hedglin et al. 2013)⍰. In this study, we analyzed sequence homology between bacterial beta clamp and archaeal/eukaryotic PCNA. Additionally, to understand their evolution, sliding clamp sequences from representative bacteria, archaea and eukarya were retrieved and phylogenetic tree was drawn for each domain of life. As expected the phylogenetic trees largely follow the vertical evolution pattern of organisms. In some instances they reveal gene duplication and horizontal gene transfer events.

## Material and methods

### Determination of sequence homology between bacterial beta clamp and archaeal PCNA

Two hundred forty nine amino acid long *Pyrococcus furiosus* PCNA sequence was obtained from Uniprot website. To discover homologues of the sequence in bacteria, the sequence was protein blasted against bacteria in the NCBI database (taxID :2). The non-redundant protein sequence database was used, and position specific iterated blast (PSI-Blast) was carried out using the default BLOSUM 62 Matrix, gap cost of Existence:11 Extension:1, with conditional compositional score matrix adjustment at the threshold of 0.005. To discover more distant homologues, second iteration of PSI-blast was carried out.

From the second iteration of the PSI blast a number of bacterial beta clamps were found as homologues of *P. furiosus* PCNA. Specifically domain 2 of bacterial beta clamp was found homologous to *P. furiosus* PCNA domain 1 and domain 3 of bacterial beta clamp was found homologous to *P. furiosus* PCNA domain 2. *Kallipyga gabonensis* beta clamp was found to be the most homologous bacterial beta clamp. To demonstrate the homology between *K. gabonensis* beta clamp and *P. furiosus* PCNA, domain 2 and 3 of *K gabonensis* beta clamp (starting amino acid 128) as discovered by pfam website was aligned with *P. furiosus* PCNA using clustal omega software. Similarly, human PCNA and *P. furiosus* PCNA, and domain 2 and 3 of *K. gabonensis* beta clamp and *E. coli* beta clamp (starting amino acid 129) were aligned using clustal omega. Earlier papers have reported that there is no homology between PCNA and beta clamp. To evaluate this point, human PCNA was aligned with domain 2 and 3 of *E. coli* beta clamp.

### Sequence Retrieval and Multiple Sequence Alignment for Tree Construction

In order to observe whether the sliding clamp proteins were conserved in their respective domain, a phylogenetic tree was constructed for different domains. The first step in this process was the sequence retrieval.

From the three domains of life, one organism each was chosen and their protein sequence for Beta-Clamp or PCNA was downloaded. The beta-clamp protein from *E*.*coli* (Uniport Accession Number : **P0A988)** was PSI-blasted against Non-Redundant Protein Sequences(NRPS) of 75 different species in bacterial domain, and all homologs of the protein from different organisms were downloaded. Similarly, the PCNA from *Pyrococcus furiosus* (Uniport Accession Number: **073947)** was PSI-blasted against Non-Redundant Protein Sequences of 74 archeal species and PCNA protein from *Homo sapiens* (**Uniport Accession Number: P12004)** was PSI-blasted against Non-Redundant Protein Sequences of 31 eukaryal species(Postberg et al. 2010)⍰ to obtain homologous proteins in each domain. The organisms in bacterial and archea domain were chosen from a list of organisms used by Moreira in 2014(Petitjean et al. 2014)^⍰^. The organisms were selected to represent all the classes, and also had their whole genome sequenced.

The obtained protein sequences were then aligned using Muscle algorithm (Edgar 2004)⍰ with default settings in MEGA X. In this way, different Multiple Sequence Alignments were generated for the three domains of life.

### Tree Construction

The evolutionary history was inferred by using the Maximum Likelihood method and JTT matrix-based model(Jones et al. 1992)^⍰^. The bootstrap consensus tree inferred from 500 replicates was taken to represent the evolutionary history of the taxa analyzed(Felsenstein 1985)^⍰^. Branches corresponding to partitions reproduced in less than 50% bootstrap replicates were collapsed. The percentage of replicate trees in which the associated taxa clustered together in the bootstrap test (500 replicates) were shown next to the branches(Felsenstein 1985)^⍰^. Initial tree(s) for the heuristic search were obtained automatically by applying Neighbor-Join and BioNJ algorithms to a matrix of pairwise distances estimated using a JTT model, and then selecting the topology with superior log likelihood value. All positions with less than 95% site coverage were eliminated, i.e., fewer than 5% alignment gaps, missing data, and ambiguous bases were allowed at any position (partial deletion option). Evolutionary analyses were conducted in MEGA X(Kumar et al. 2018)^⍰^.

## Result and discussion

### Detection of homology between PCNA and sliding clamp

With a threshold of 0.005, *P. furiosus* PCNA was PSI blasted against the NCBI non-redundant bacterial protein sequence database to the second iteration. Figure 1A shows a screenshot of the top hits from the blast. A number of PCNAs were discovered in the bacterial genome. Given that PCNAs are characteristic proteins of eukaryotes and archaea, these proteins may have arisen by horizontal gene transfer from eukaryotes or archaea to bacteria. Further analysis is necessary to determine their source of origin. The proteins not found in the first iteration, but found in the second iteration are highlighted in yellow as in Figure 1b. A slew of bacterial beta clamp were discovered as homologues of *P. furiosus* PCNA upon PSI blast at second iteration. Top among the hits was *Kallipyga gabonensis* beta clamp. The query cover for the blast was 94 percent, E score was 1 × 10^−4^, and sequence identity was above 19%. Since a random blast of PCNA yielded beta clamp of various species, it can be claimed that the two are homologous.

**Fig 1.**
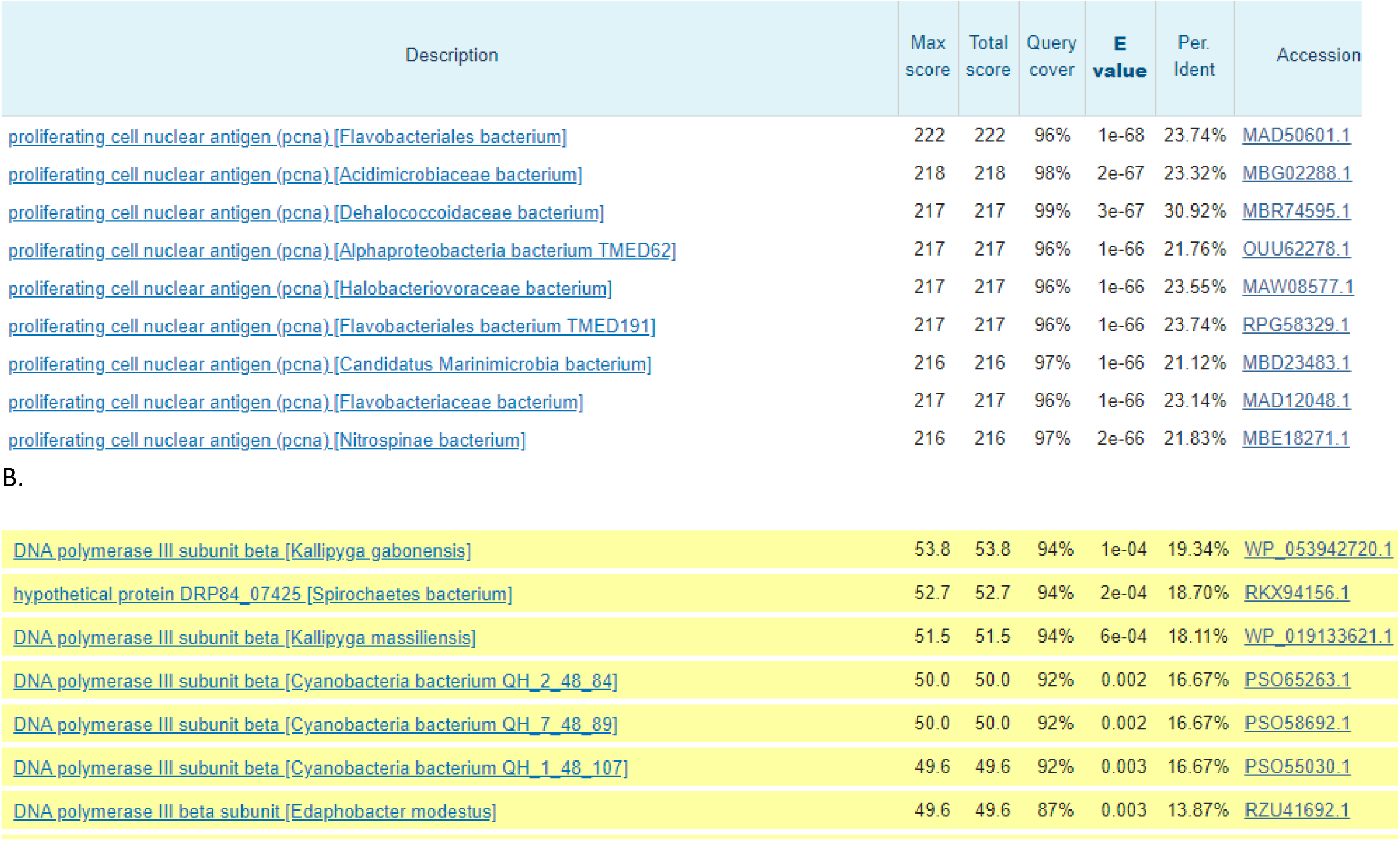
Screenshot of the results obtained when *P. furiosus* PCNA is PSI blasted against the NCBI bacterial database. (A) shows the results obtained when regular blast is carried out. PCNA sequences in undefined bacterial genomes appear. It is possible that these bacterial PCNAs emerged in bacterial genome from horizontal gene transfer from archaea and eukarya. (B) shows bacterial beta clamp from some defined and some undefined species that appear in the blast after second iteration. The yellow color represents hits that only appear in the second iteration of the blast. *Kallipyga gabonensis* is the top defined beta clamp hit in the bacteria. Other defined bacterial beta clamps that appear in the blast search are *Kallipyga massiliensis* and *Edaphobacter modestus* beta clamps. Query cover above 90 percent in these searches indicates that the entire PCNA was covered during the blast search.

**Fig 2.**
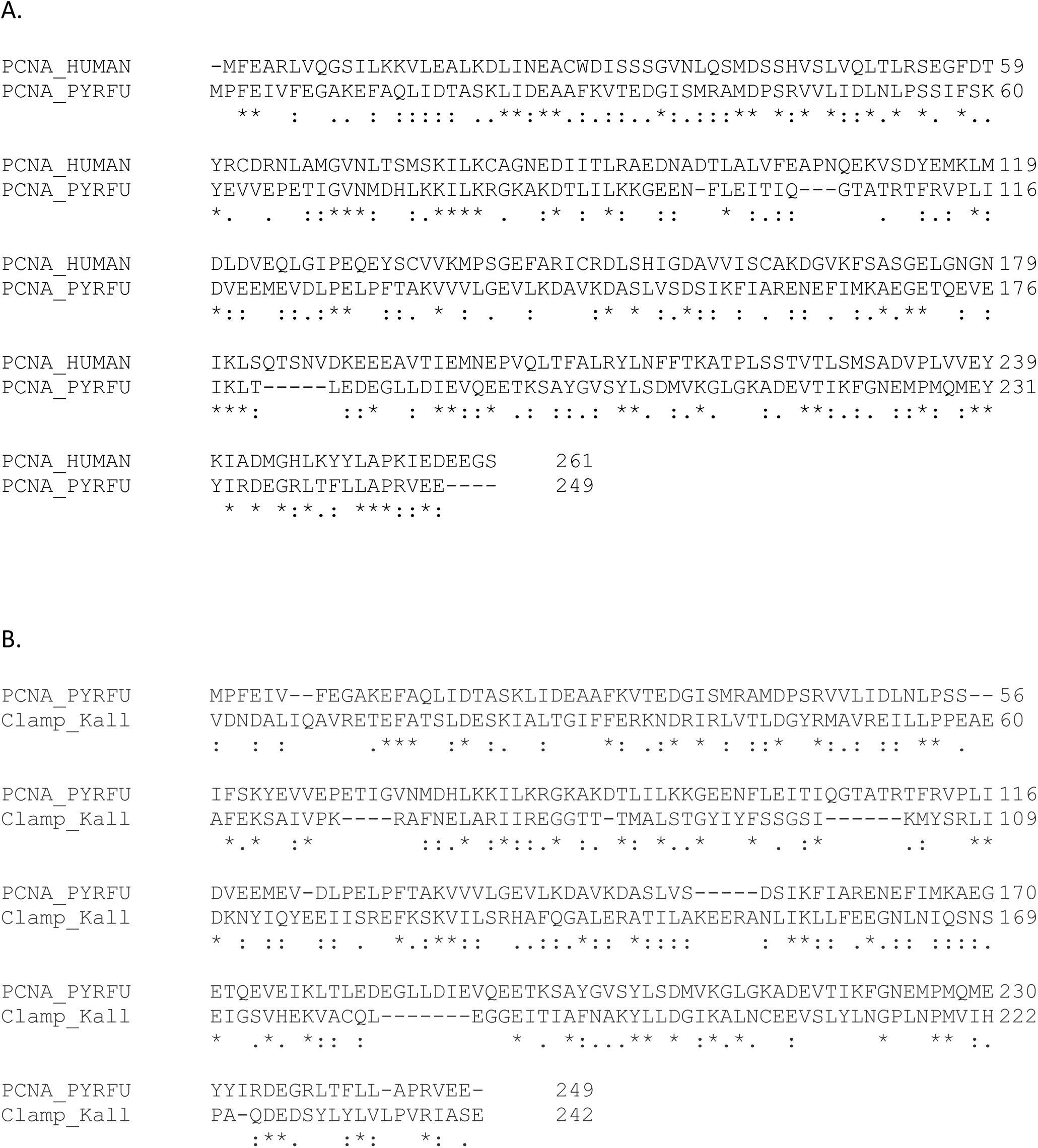
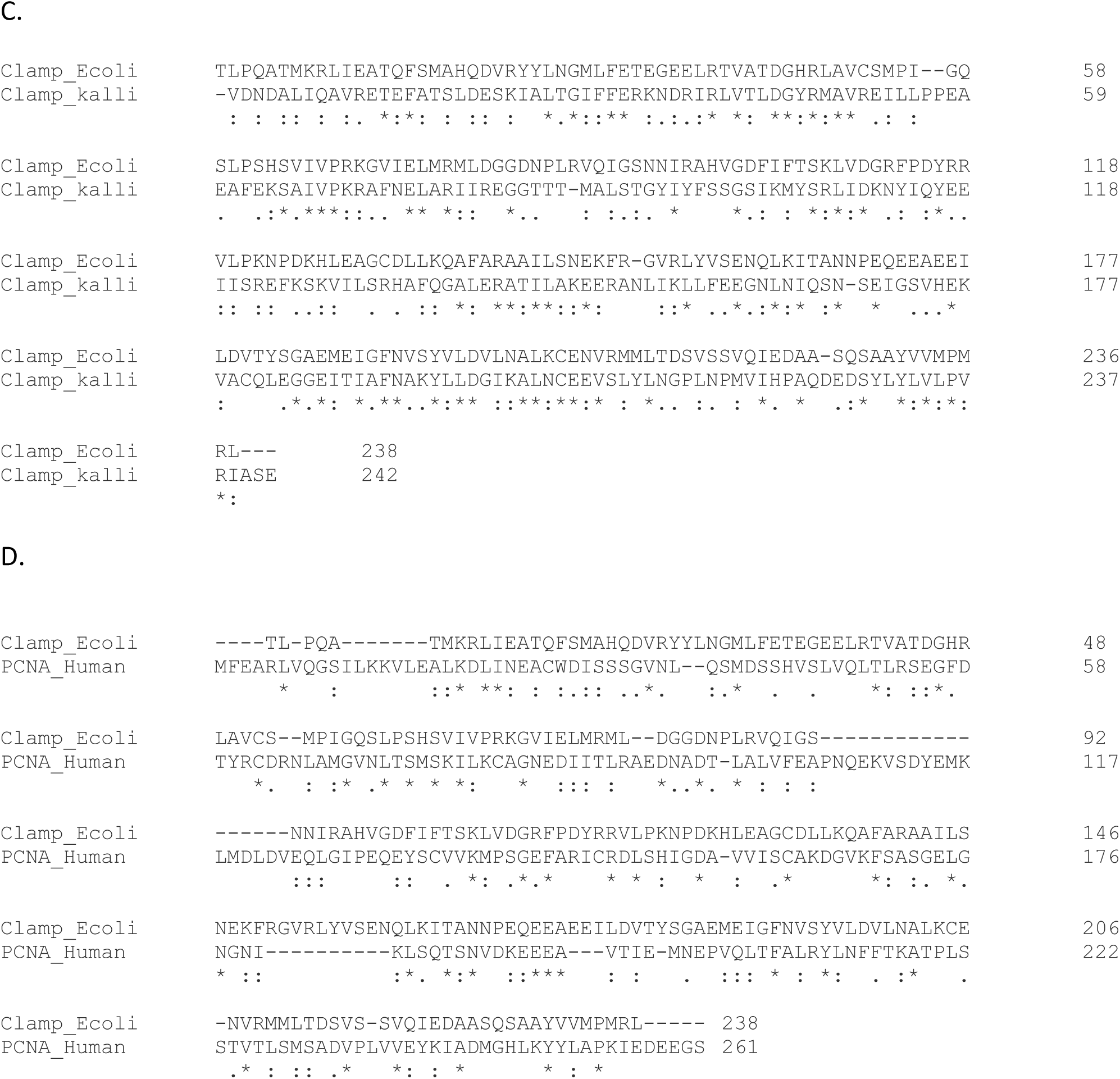
Clustal omega alignment of (A) Human and *P. furiosus* PCNAs (B) *P. furiosus* PCNA and *K. gabonensis* beta clamp (C) *E. coli* beta clamp and *K. gabonensis* beta clamp (D) *E coli* beta clamp and human PCNA. When PCNA and beta clamps were aligned, domain 1 of PCNA was aligned to domain 2 of beta clamp and domain 2 of PCNA was aligned to domain 3 of beta clamp. Since (A) and (C) align PCNA to PCNA and beta clamp to beta clamp (domains 2 and 3 aligned), they show the highest degree of similarity and identity. (B) *P. furiosus* PCNA and *K. gabonensis* beta clamp also display high degree of sequence identity and sequence similarity that is statistically significant. In the figure * represents sequence identity, : represents high sequence similarity (similar category of amino acids) and. represents low sequence similarity.

A closer inspection of the alignment result shows that out of the three domains of beta clamp, domains 2 and 3 align with the full length of the PCNA. This suggests that either the Lowest Universal Common Ancestor (LUCA) had three domains in its sliding clamp, the first of which got lost in archaea, or LUCA had two domains in its sliding clamp and the third domain got added after duplication event. The alignment of *K. gabonesis* beta clamp and *P. furiosus* PCNA shows 20%(50/249) sequence identity, 44%(109/249) sequence similarity and 11%(27/249) gaps. On the other hand, *P. furiosus* PCNA and human PCNA show 25%(61/249) sequence identity, 61%(136/249) sequence similarity and 5%(13/249) gaps. *E. coli* beta clamp and human PCNA, which were earlier claimed to not have sequence homology, show 16%(42/261) sequence identity, 36%(94/261) sequence similarity and 16%(43/261) gap. These results demonstrate that although more common model organism (human and *E. coli*) PCNA and beta clamp do not demonstrate high degree of homology, non-model organism (*K. gabonesis* and *P. furiosus*) beta clamp and PCNA demonstrate statistically noteworthy sequence homology.

Given that domain 2 and 3 of beta clamp align with the two domains of PCNA, it can be hypothesized that domain 1 from beta clamp interacts with bacteria specific replication and repair protein. Most of the interaction assays carried out with PCNA and beta clamp have been conducted with the entire protein. It would be interesting to zero in on which domain of the sliding clamps is important for individual interactions.

### Phylogenetic tree of eukaryotic PCNA

To study the origin and evolution of eukaryotic PCNA, PCNA from human was protein blasted in NCBI against the non-redundant protein sequences of selected eukaryotic organisms from different classes. Certain species of the kingdom/class eozoa, amoebozoa, ciliophora, archaeplastida, fungi and animalia were chosen. Where multiple PCNAs were discovered all sequences were downloaded. A phylogenetic maximum likelihood tree was drawn using these sequences. A tree thus constructed is shown in Figure 3. The proteins from different classes tended to segregate as labeled on the right. For example all PCNAs from ciliophora and animalia cluster together. This demonstrates that there is very little horizontal gene transfer. A largely vertical inheritance pattern can be expected for an essential protein like PCNA. However, a number of species in ciliophora class demonstrate gene duplication events. Three PCNA genes are found in both *Stylonychia lemnae* and *Oxytricha trifallax*. Each of the three proteins from one species lies next to a protein from another species. This suggests that a single PCNA gene underwent gene duplication twice in the progenitor organism, which gave rise to three PCNAs, each of which evolved separately when speciation occurred.

**Fig 3.**
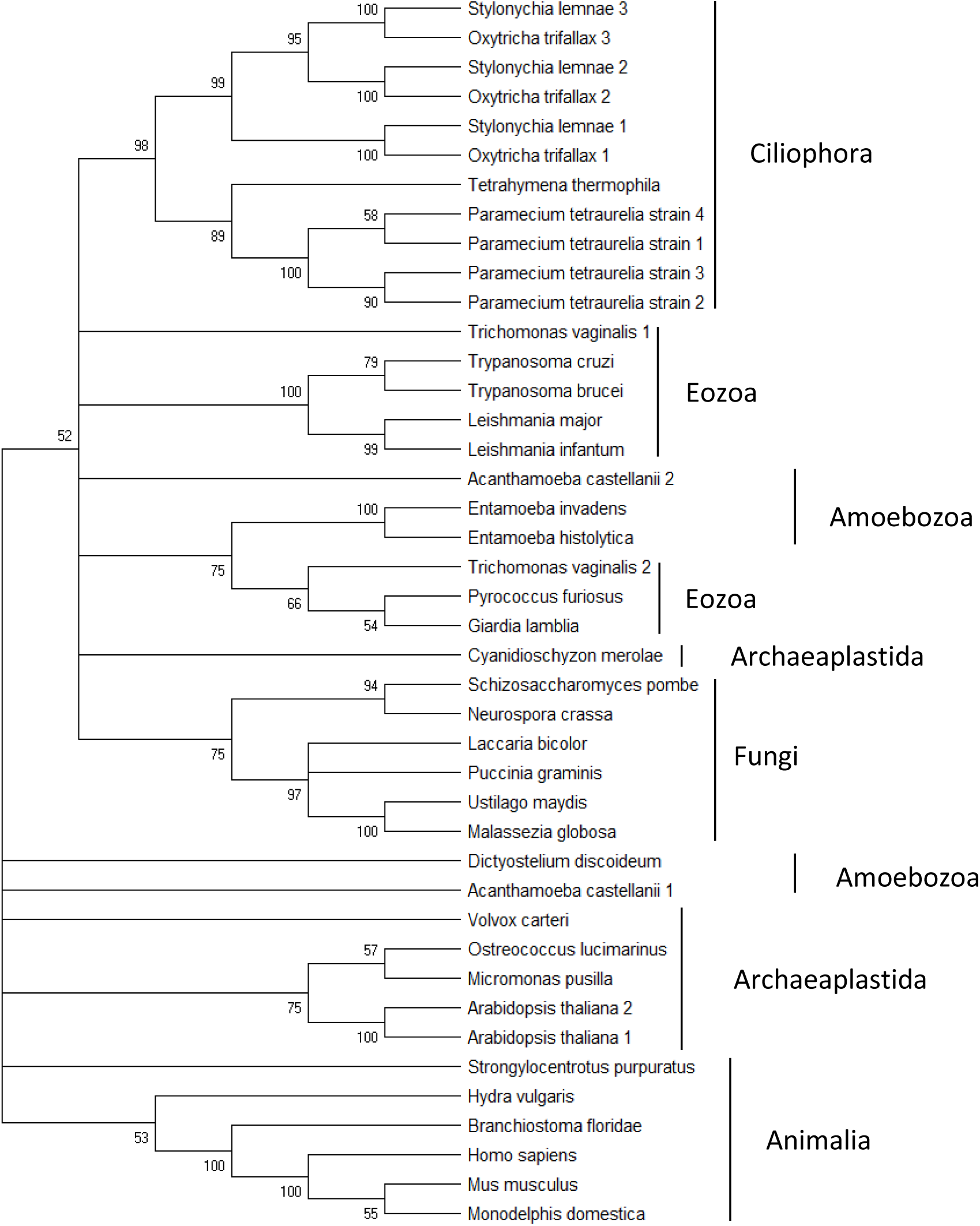
Phylogenetic Maximum Likelihood tree drawn for PCNA of 31 different eukaryotic species. Both domains of PCNA were used for alignment. Species from 6 kingdoms of eukaryotes—animalia, fungi, archaeplastida, amoebozoa, eozoa and ciliophora (inside the kingdom chromista)—were used to draw the tree. The purpose of using organisms from 6 different kingdoms is to represent the entire eukaryotic domain, not just the crown groups. This tree demonstrates that most species contain only one PCNA that is vertically inherited. The instances where there is gene duplication and horizontal gene transfer is indicated in the text.

The figure also demonstrates that there are four closely related PCNAs in *Paramecium tetraurelia*. The tree pattern suggests that there was an initial duplication event, followed by duplications of the duplicated PCNA.

Amoebozoa and eozoa PCNAs tended to cluster at two locations each. *Trichomonas vaginalis* had two PCNAs, one clustering with other eozoa PCNAs and other clustering with *Pyrococcus furiosa* (archaea) PCNA. This demonstrates that one PCNA was vertically inherited, while other PCNA might have arisen from horizontal gene transfer from archaea. Other eozoa (*Leishmania* and *Trypanosoma*) PCNA clustered together, suggesting that they all came from a common ancestor. *Giardia lambia* (an eozoa) PCNA, however segregated separately close to *Pyrococcus furiosus* PCNA, hinting at a possible archaeal origin.

As for amoebozoa PCNAs, three PCNAs from different species lie next to each other, whereas two other PCNAs, one from *Dictostylium discoideum* and another from *Acanthamoeba castellani*, are sandwiched between archeaplastida and fungi. It has been observed from earlier research that *Dictostylium* molecules resemble archeaplastida and fungi molecules more than amoebozoa molecules(Baldauf & Doolittle 1997; Eichinger et al. 2005)□. So this placement in the tree does not come as a surprise. The second *Acanthamoeba castellani* PCNA might have arisen from horizontal gene transfer from *Dictosylium* or archaeaplastida or fungi.

All fungi and animalia PCNAs cluster together suggesting a common origin. There seems to be no gene duplication event in the species we have considered in the paper. All except one archaeplastida molecules cluster together also suggesting a common origin. *Cyanidioschyzon merolea* PCNA is found at a different location from other archaeplastida genome suggesting a different ancestry.

### Phylogenetic tree of bacterial beta clamp

The Beta-clamp from *E. coli* was PSI-blasted against non-redundant protein sequences from different organisms belonging to different classes in bacteria. A total of 85 homologs of the protein were identified from 75 species of bacteria. All the homologs were downloaded, and aligned together to build a phylogenetic tree as shown in figure 4. It was observed that most of the bacteria consisted of only one homolog for Beta-clamp. However, some bacteria had two or more homologs of the protein, suggesting ancient gene duplication or horizontal gene transfer. The bacteria belonging to same phylum, or sharing similar properties were found to be clustered together apart from some exceptions, which suggests that the beta-clamp protein is conserved among similar species in bacterial domain.

**Fig. 4.**
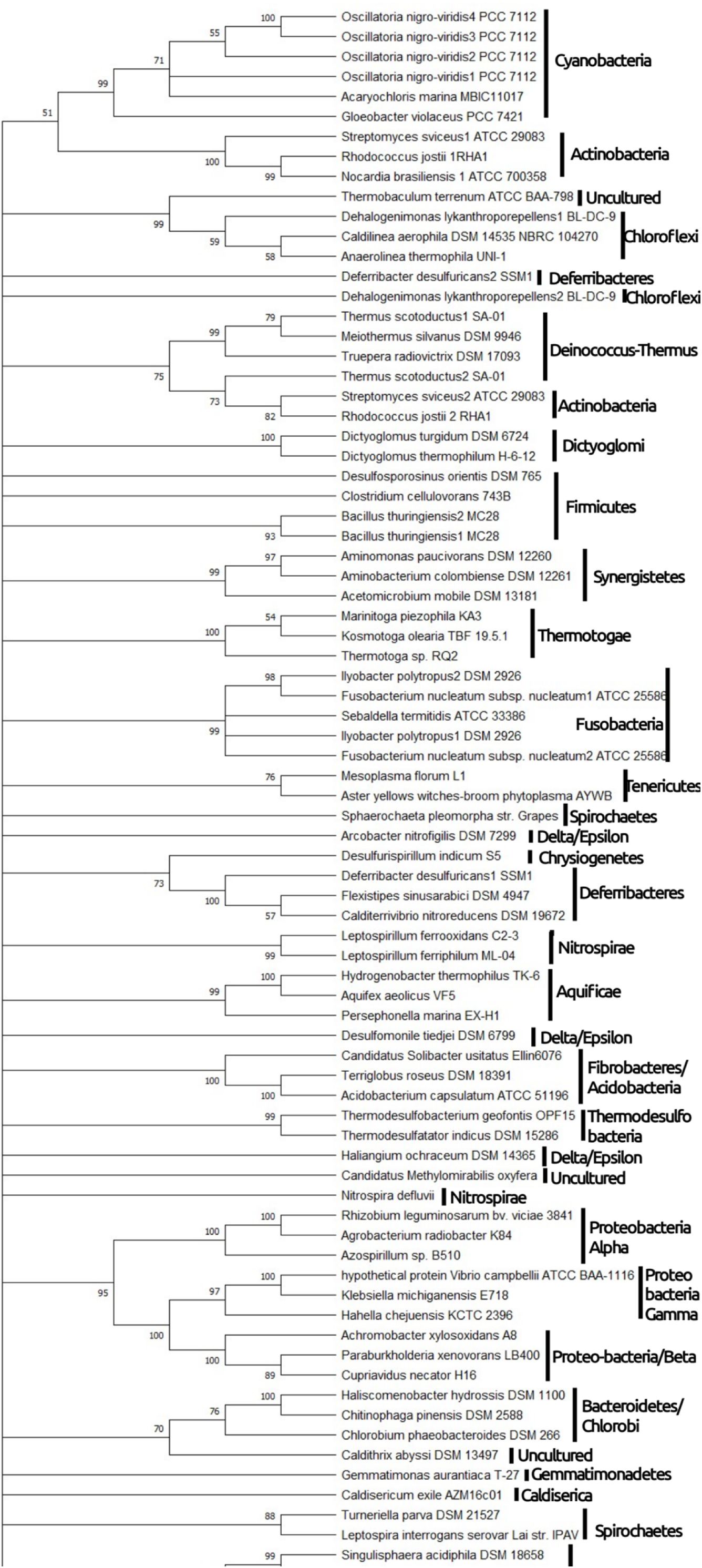
Maximum Likelihood tree drawn for beta-clamp of 75 different bacterial species representing different phylum and classes. All three domains of beta-clamp were used for alignment. The tree demonstrates that most bacteria have only a single homolog for beta-clamp protein, which is vertically inherited. Some instances of gene duplication and horizontal gene transfer are also observed. Bacteria belonging to the same class are clustered together with minor exceptions.

In proteobacteria, three classes viz. Alphaproteobacteria, Betaproteobacteria and Gammaproterbacteria are found to be clustered together, originating from a single node suggesting common origin. However, the Delta/Epsilon sub-divisions seem to scatter away from other Proteobacteria in the phylogenetic tree. It suggests that the the phylum Proteobacteria is not monophyletic which has also been previously suggested(Hug et al. 2016)□. The phyla Planctomycetes, Verrucomicrobia and Chlamydiae which are often grouped together as PVC superphylum(Hug et al. 2016)^□^ were observed to be clustered together, suggesting similar origin and properties.

Multiple homologs of the same protein in a species can be accounted for by ancient gene duplication or horizontal gene transfer events. In case of *Oscillatoria nigroviridis*, four homologs of sliding beta clamp were observed. It was a result of three ancient gene duplication at different stages of time, followed by vertical gene transfer. Similar was the case for *Bacillus thirungenesis*. The resulting homologs didn’t undergo much variation and thus were clustered together. The phylum Actinobacteria was positioned at two different locations in the phylogenetic tree, one clustered with Cyanobacteria and the other with Deinococcus-thermus. It suggests that the horizontal gene transfer occurred from the ancestor of Deinococcus-thermus to actinobacteria before speciation. This gene was maintained in some species, which showed two homologs for beta-clamp protein, while disappeared in some species (*Nocardia brasiliensis*), which had only a single homolog.

In the phylum Fusobacteria, two homologs each of *Ilyobacter polytropus* and *Fusobacterium nucleatum subsp. nucleatum* were observed, which were clustered together. It might have resulted from ancient gene duplication, or lateral gene transfer among the species. Two homologs of the beta-clamp protein of *Deferribacter desulfuricans* were observed, where one homolog was separated from its counterpart and located between species of Chloroflexi suggesting lateral gene transfer from Chloroflexi. Other cases of lateral gene transfer can be claimed in *Nitrospira defluvii*, and *Dehalogenimonas lykantroporepellens*.

Apart from some cases of horizontal gene transfer and gene duplication, the species from same class were clustered together in the phylogenetic tree. Thus, we can conclude that beta-clamp protein is fairly conserved in bacterial domain.

### Phylogenetic Tree of Archeal PCNA

The PCNA from *P. furiosus* was PSI-blasted against non-redundant protein sequences from different archaea belonging to different classes in archea. 115 homologs of PCNA were identified in 74 species of archaea. All of the homolog sequences were downloaded, and a multiple sequence alignment and phylogenetic tree was constructed similarly as discussed above.

The obtained phylogenetic tree is shown in Figure 5. It can be observed that the three main phyla of domain archea, viz Thaumarchaeota, Euryarchaeota and Crenarchaeota were positioned separately in the phylogenetic tree, apart from some exceptions. The phylum Thaumarchaeota was sandwiched in between Euryarchaeota and Crenarchaeota. A single species of Korachaeota was present within the phylum Crenarchaeota, which leads to questions the monophyly of Crenarchaeota.

**Fig. 5.**
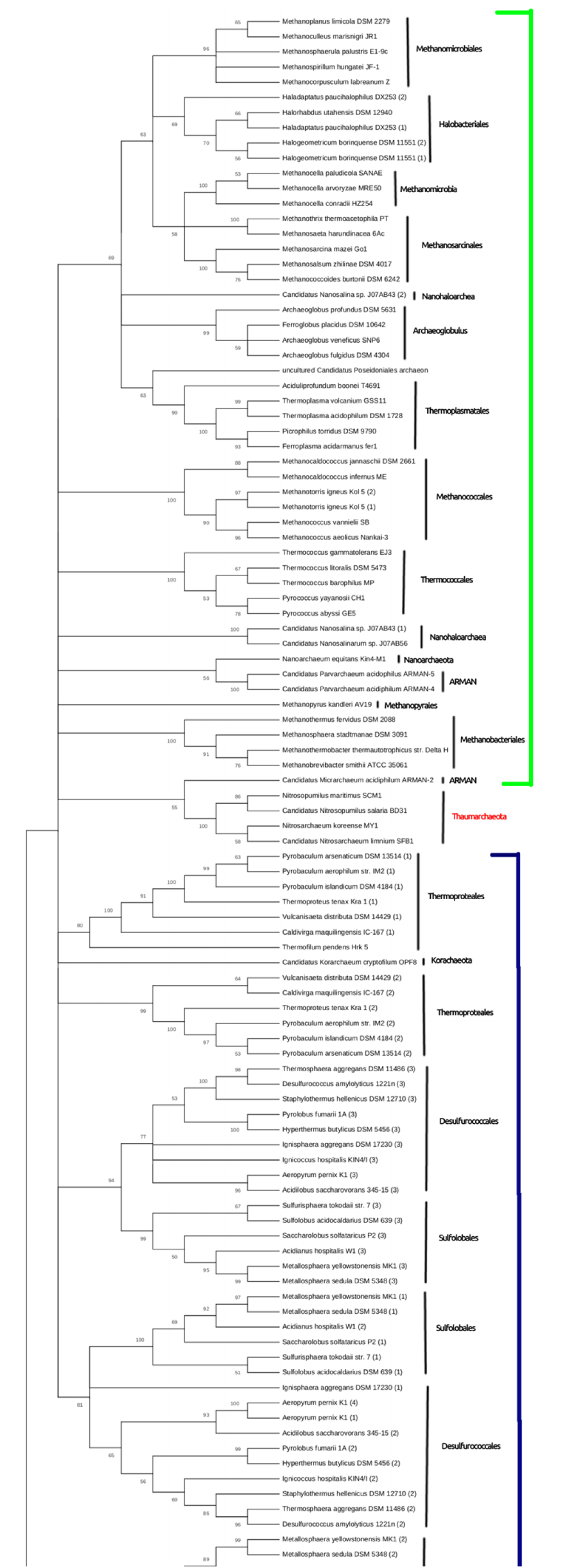
Maximum Likelihood tree drawn for PCNA of 75 different archaeal species representing three major phyla: Euryarchaeota, Thaumarchaeota and Crenarchaeota (Green Line: Euryarchaeota, Blue Line: Crenearchaeota). The tree demonstrates that the three phyla are positioned separately. Some archea are observed to have more than one homolog for PCNA. It is observed that PCNA protein is highly conserved in archaea, with few exceptions of gene duplication and horizontal gene transfer.

In the phylum, Euryarchaeota, the species belonging to the same class were classified together. However, some cases of duplication and horizontal gene transfer were observed. Two homologs of PCNA were observed in *Halogeometricum borinquense DSM 11551*, which were separated from the same node, hence suggesting ancient gene duplication. Similar was the case for *Methanotorris igneus Kol5*. Also, two homologs of the protein were discovered for *Haladaptatus paucihalophilus*, and *Canditatus Nanosalina*. These two homologs were not positioned together, hence suggests horizontal gene transfer.

The phylum Crenarchaeota is divided into three classes, viz. Sulfolobales, Desulfurococcales, and Thermoproteales. The phylogenetic tree obtained still supports the close relationship of Sulfolobales and Desulfurococcales, also mentioned by Armanet(Brochier-Armanet et al. 2011)^□^□. Both these classes had three homologs of PCNA. The two homologs appear to have originated by gene duplication in the ancestor of these two classes, followed by modification to form different proteins, while the third homolog might have been acquired by lateral gene transfer from other diverge species, possibly eukaryotes. As all three homologs are required for the assembly of a functional hetero-trimer, the homologs are observed to be vertically inherited in future generations. However, the third phylum of Crenarchaeota: Thermproteales had two homologs of PCNA. Thus, it can be predicted that the functional PCNA in Thermoproteales is a homo-trimer (of any one of the homolog) and two homologs might have arisen due to ancient gene duplication.

The phylum Euryarchaeota can be divided into two groups: Sub-Phyla I and Sub-Phyla II. Sub-Phyla I consists of Thermcoccales, Nanoarchea, and class I methanogens, while Sub-Phyla II consists of Archeoglobulus, Halobacteriales, Thermoplasmatales and Class II methanogens.(Forterre 2015)^□^ The phylogenetic tree in figure 5 suggests the monophyly of Sub-Phyla II, which is also supported by the updated tree of life by Forteree.(Forterre 2015) □ However, the monoplyly of Sub-Phyla I couldn’t be supported statistically.

The phylogenetic tree further clarifies the position of Methanopyrales in the phylogenetic tree. As *Methanopyrales kandleri* was positioned along with Methanobacteriales, Methanopyrales can be classified as Class I Methanogen. However, the monophyly of Class I and Class II methanogen couldn’t be supported. Methanobacteria, Methanobacteriales and Methanosarcinales were observed to share a common ancestor along with Halobacteriales. Methanococcus was positioned near to Thermococcales, while Nanohaloarchea and Nanoarchaeota were sandwiched between Methanococcus and Methanobacteriales.

The phyum Nanoarchaeota was positioned near to Thermococcales, along with Nanohaloarchaea, and ARMAN group. The close association of Nanoarchaeota to Thermococcales was also observed by Brochier in 2005(Brochier et al. 2005)^□^. Also, Thermococcales, Nanoarchaea, ARMAN and Nanohaloarchea are placed together suggesting they are closely related. It has also been supported by phylogeny based on ribosomal proteins and DNA replication proteins(Raymann et al. 2014; Forterre 2015)^□^. The tree however, questions the position of ARMAN-2. It has been positioned as a sister group to Thaumarchaeota, forming a border-line between Crenearchaeota and Euryarchaeota.

Thus, the phylogenetic tree based on archaeal PCNA follows vertical pattern of evolution in most cases. Also, archael PCNA is conserved among closely related species of the Archaea domain.

To conclude, the sliding clamp proteins (Beta-Clamp in Bacteria, PCNA in Archaea and Eukarya) are highly conserved within their respective domain apart from some exceptions. Also, the observed sequence homology between some Beta-Clamps and PCNA generates the possibility of common origin for the sliding clamp proteins.

